# Peds1 deficiency in zebrafish results in myeloid cell apoptosis and exacerbated inflammation

**DOI:** 10.1101/2023.09.26.559500

**Authors:** Ana B. Arroyo, Sylwia D. Tyrkalska, Eva Bastida-Martínez, Antonio J. Monera-Girona, Joaquín Cantón-Sandoval, Martín Bernal-Carrión, Diana García-Moreno, Montserrat Elías-Arnanz, Victoriano Mulero

## Abstract

Plasmalogens are glycerophospholipids with a vinyl ether bond that confers unique properties. Recent identification of the gene encoding PEDS1, the desaturase generating the vinyl ether bond, enables evaluation of the role of plasmalogens in health and disease. Here, we report that Peds1-deficient zebrafish larvae display delayed development, increased basal inflammation, normal hematopoietic stem and progenitor cell emergence, and cell-autonomous myeloid cell apoptosis. In a sterile acute inflammation model, Peds1-deficient larvae exhibited impaired inflammation resolution and tissue regeneration, increased interleukin-1β and NF-κB activities, and elevated ROS levels at the wound site. Abnormal immune cell recruitment, neutrophil persistence, and fewer but predominantly pro-inflammatory macrophages was observed. Chronic skin inflammation worsened in Peds1-deficient larvae but was mitigated by exogenous plasmalogen, which also alleviated hyper-susceptibility to bacterial infection, as did pharmacological inhibition of caspase-3 and colony-stimulating factor 3-induced myelopoiesis. Overall, our results highlight an important role for plasmalogens in myeloid cell biology and inflammation.

**Keypoints:**

- Plasmalogens are crucial for cell autonomous survival, recruitment and activation of neutrophils and macrophages.
- Plasmalogen production aids inflammation resolution, while supplementation reduces inflammation and boosts bacterial clearance.

## Introduction

The function and physicochemical properties of cellular membranes are predominantly determined by the chemical and compositional diversity of lipids. Alterations in membrane lipid homeostasis are related to numerous diseases, but the causality and the underlying molecular mechanisms are often unclear^1^. Glycerophospholipids (GPs) are the major membrane lipids and consist of a glycerol backbone with fatty acids at the *sn*-1 and *sn*-2 positions, typically bound via an ester bond. However, a significant subset of GPs, generally known as ether lipids, contain an ether-linked fatty alcohol side chain at the *sn*-1 position. These comprise plasmanyl GPs, with an *sn*-1 alkyl group (1-*O*-alkyl, linked by an ether bond) and plasmenyl GPs, with an *sn*-1 alkenyl group (1-*O*-alk-1′-enyl, with a cis-double bond conjugated with the ether oxygen or vinyl ether bond). The latter are called plasmalogens and are the most abundant subgroup of ether lipids.

Ether lipids have a unique distribution across the evolutionary tree, as they are found in strictly anaerobic bacteria, protozoans, and metazoans, where they comprise 5-20 % of total GPs content, but are absent in fungi, plants and most aerobic bacteria, except myxobacteria. Ether lipid composition varies among organisms, organelles, and cell types. In mammals, they are abundant in the brain, heart, kidney, lung, skeletal muscle, and neutrophils, but are scarce in the liver. They are present in nearly all subcellular membranes — plasma membrane, lipid droplets, mitochondria, endoplasmic reticulum, and post-Golgi network — except for peroxisomes, where the early biosynthetic steps occur^2,3^. Formation of the first intermediate with an ether bond (1-*O*-alkyl-glycerone phosphate) in the peroxisome requires three enzymes: GNPAT (glycerone phosphate *O*-acyltransferase), FAR1/FAR2 (fatty acyl-CoA reductase), and AGPS (alkylglycerone-phosphate synthase). Four more steps in the endoplasmic reticulum lead to the synthesis of plasmanylethanolamine, which is the substrate of PEDS1 (plasmanylethanolamine desaturase 1), an enzyme whose identity as the product of the *TMEM189* gene was only recently discovered ^3-6^. PEDS1 generates the vinyl ether bond, producing ethanolamine plasmalogens (PE-PLs), which are converted to the choline form (PC-PLs) in subsequent steps. These are highly enriched in the heart and smooth muscle, while PE-PLs are the most abundant form in the rest of the organs ^7^.

Besides differing from their ester counterparts, alkyl and alkenyl ether lipids also have different physicochemical properties and, thus, may have different roles in cell physiology. These differences affect membrane packing, curvature and fluidity, as well as interaction with other lipids ^8^. Plasmalogens are also proposed to act as molecular scavengers that protect membranes from oxidative damage, since the vinyl ether bond is highly susceptible to cleavage by reactive oxygen species (ROS), yielding products with signaling functions ^2^. Until recently, most studies could not discern the specific biological roles of plasmalogens from those of their alkyl ether lipid precursors. However, the discovery of PEDS1 identity has spurred investigation about the exact functions of plasmalogens and the molecular basis of their actions. This orphan enzyme was first identified from delving into the photooxidative stress response of the soil myxobacterium *Myxococcus xanthus.* The study revealed that CarF and its homologs in animals (human, mouse, worm, fly and the two zebrafish paralogs), but not those in plants, encode the long-sought PEDS1 ^6^. Soon after, two other studies identified the desaturase for human plasmalogen biosynthesis by completely orthogonal approaches ^4,5^.

PEDS1 has been recently involved in cell fitness under hypoxia conditions ^9^ and in negative regulation of autophagy ^10^. A context-dependent role of plasmalogens in ferroptosis has been reported, since some studies find PEDS1 depletion to have no effect on ferroptosis ^11,12^, while others suggest a pro-ferroptotic effect ^13,14^. *In vivo* and *in vitro* experiments suggest that PEDS1 deficiency reduces progression of gastric and breast cancer ^10,14^. The PEDS1-knockout mouse has several associated phenotypes, such as decreased growth, altered blood parameters and eye abnormalities ^15,16^. However, the molecular basis of these phenotypes remains elusive.

The relevance of ether lipids is highlighted by their involvements in several human pathologies: rare genetic disorders associated with mutations in peroxisomal enzymes involved in ether lipid biosynthesis (Zellweger syndrome and Rhizomelic chondrodysplasia punctata-RCDP) or complex diseases, such as Alzheimer’s and Parkinson’s diseases (AD and PD, respectively), multiple sclerosis, autism, schizophrenia, psychiatric depression, and metabolic and inflammatory disorders, including diabetes mellitus, cancer and various respiratory and cardiac diseases. These data suggest that slight differences in lipid chemical structures are critical for cellular functions ^17^.

Plasmalogens appear to have a dual role in inflammation. On the one hand, the *sn*-2 position of plasmalogens is mostly occupied by PUFAs (fatty acids with more than one double bond), such as arachidonic acid, which serves as a precursor to produce pro-inflammatory lipid mediators. However, the lack of plasmalogens does not necessarily cause a deficit in PUFAs and, thus, they are not essential reservoirs of these lipid mediators ^18^. On the other hand, plasmalogens may have anti-inflammatory functions, since chronic inflammation is a common element in pathophysiological conditions with decreased plasmalogens levels. In fact, plasmalogen replacement therapy (PRT), based on administration of plasmalogens or their precursors to normalize plasmalogens levels, led to promising results in several disease models studies. PRT not only improved phenotypes of patient-derived cells and animal models with peroxisomal disorders and neurodegenerative diseases, but it was also reported to ameliorate symptoms in patients with AD or PD ^7,19^. Despite the observed interrelationship, it is less well understood whether the decrease of plasmalogens is either the cause or the consequence of chronic inflammation-linked diseases.

In the present study, we used the unique advantages of the zebrafish to unequivocally show the relevance of plasmalogens in inflammation. Thus, Peds1 deficiency resulted in cell-autonomous neutrophil and macrophage apoptosis and, concomitantly, exacerbated inflammation. Moreover, in a sterile model of acute inflammation, Peds1 deficiency provoked defective inflammation resolution and tissue regeneration, accompanied by neutrophil persistence and predominantly pro-inflammatory macrophages at the injury site. Additionally, Peds1 deficiency also aggravated chronic skin inflammation and caused hyper-susceptibility to bacterial infection.

## METHODS

### Animals and ethical statement

Zebrafish (*Danio rerio* H.) were obtained from the Zebrafish International Resource Center (ZIR) and mated, staged, raised and processed according to the zebrafish handbook ^20^. The lines *Tg(mpx:eGFP)^i114^* ^21^, *Tg(mpx:Gal4.VP16)^i222^*^22^, *Tg(lyz:DsRED2)^nz50^* ^23^, *Tg(mfap4.1:mCherry-F)^xt12^*^24^, *Tg(gata1a:DsRed)^sd2^*and *Tg(-6.0itga2b:eGFP)^la2^* ^25^, *Tg(NFkB-RE:eGFP)^sh235^*^26^, *Tg(il1b:GFP-F)^ump3^* ^27^, *Tg(mfap4.1:mCherry-F), Tg(tnfa:eGFP-F)^ump5^* ^28^, *mitfa^w2/w2^; mpv17^a9/a9^*, referred to here as casper, ^29^, and *spint1a^hi2217Tg/hi2217Tg^*^30^ were previously described. The experiments performed comply with the Guidelines of the European Union Council (Directive 2010/63/EU) and the Spanish RD 53/2013. The experiments and procedures performed were approved by the Bioethical Committees of the University of Murcia (approval number #669/2020).

### Gain- and loss-of-functions experiments

Crispr RNA (crRNA) for zebrafish genes *peds1a and peds1b*, *il1b* and negative control **(Table S1),** and tracrRNA were purchased from Integrated DNA Technologies (IDT) and resuspended in nuclease-free duplex buffer (DB) up to 100 μM. For duplexing 1μl of each was mixed and incubated for 5 min at 95°C. After cooling down to room temperature (RT), the duplex was diluted to 1000 ng/µl by adding 1.43 µl of Nuclease-Free Duplex Buffer. Finally, the injection mix was prepared by mixing 1 μl of duplex, 2.55 μl of DB, 0.25 μl Cas9 nuclease V3 (IDT, #1081058) and 0.25 μl of phenol red, giving final concentrations of 250 ng/μl of gRNA duplex and 500 ng/μl of Cas9. The prepared mix was microinjected into the yolk of one-to eight-cell-stage embryos using a microinjector (Narishige) (0.5-1 nl per embryo). For checking the efficiency of crRNA, genomic DNA from a pool of 10 microinjected larvae was extracted with the HotSHOT method ^31^ and used as template to amplify the target sequences with a specific set of primers **(Table S1)**. Sanger sequencing data were analyzed with the TIDE webtoolM (https://tide.nki.nl/) and/or SYNTHEGO Crisper Performance Analysis webtool (https://ice.synthego.com).

In vitro-transcribed RNA (*csf3a*) was obtained following manufacturer’s instructions (mMESSAGE mMACHINE kit, Ambion) from a *csf3a* construct described previously ^32^. RNA was mixed in microinjection buffer (×0.5 Tango buffer and 0.05% phenol red solution) and microinjected into the yolk sac of one-cell-stage embryos using a microinjector (Narishige; 0.5–1 nl per embryo). The same amount of RNA was used in all experimental groups.

The *uas:peds1; mylf7:eGFP* construct was generated by MultiSite Gateway assemblies using LR Clonase II Plus (ThermoFisher Scientific) according to standard protocols and using Tol2 kit vectors described previously ^33^. The *uas:peds1; mylf7:eGFP* construct (40 ng/µl) was microinjected (0.5-1 nl) into the yolk sac of one-cell-stage *Tg(mpx:Gal4.VP16)^i222^* embryos together with Tol2 RNA (155 ng/μl) in microinjection buffer. Embryos were sorted at 2 dpf under a fluorescence stereomicroscope using the heart fluorescent marker of the construct.

### Lipid extraction and GC-MS analyses

Each sample of approximately 50 larvae was collected by centrifugation, washed once with cold PBS and stored at -80 °C until further use. Lipid extraction and fatty acid methyl ester (FAME) analysis were performed following a previously described protocol with slight changes (Gallego-García et al., 2019). In brief, the pellets were transferred to Pyrex glass tubes, resuspended in 4 ml of chloroform:methanol (2:1 v/v) by vortexing and incubated for 1 hour in ice. Then, they were mixed with 1 ml of 120 mM KCl, vortexed for 10 seconds and centrifuged at 400 g for 5 minutes at 4 °C to induce phase separation. After removing the upper aqueous/protein phase, the lower phase was passed through a Whatman No. 1 filter paper into a new Pyrex glass tube and dried under a N_2_ stream at 35°C. For FAMEs generation, dried samples were subjected to acid hydrolysis at 55 °C overnight in 0.5 ml toluene and 1 ml of 1% H_2_SO_4_ in methanol. The reaction was quenched with 1 ml of 0.2 M KHCO_3_, mixed gently with 2.5 ml hexane:diethyl ether (1:1 v/v) containing 0.01% 2,6-di-tert-butyl-4-methylphenol (BHT; Scharlab) to minimize oxidation and centrifuged at 500 g for 2 minutes at 4 °C. The upper hexane:ether phase was transferred to a separate tube and the lower phase was re-extracted with 2.5 ml hexane:diethylether (1:1 v/v). The combined upper phase was dried with N_2_ at 35 °C and resuspended in 200 µl hexane. Alternatively, to also analyse ether lipid-derived OAGs, dried samples were resuspended in 162.5 µl hexane and 37.5 µl N-methyl-N-trimethylsilyltrifluoroacetamide (MSTFA) (Sigma-Aldrich) and incubated at 37 °C for 1 h. GC-MS and subsequent data analyses were performed as indicated previously (Gallego-García et al., 2019).

### Analysis of gene expression

Total RNA was extracted from the tail part of the zebrafish body with TRIzol reagent (Invitrogen) following the manufacturer’s instructions and treated with DNase I, amplification grade (1 U/mg RNA: Invitrogen). SuperScript IV RNase H Reverse Transcriptase (Invitrogen) was used to synthesize first-strand cDNA with random primer from 1mg of total RNA at 50 °C for 50 min. Real-time PCR was performed with an ABIPRISM 7500 instrument (Applied Biosystems) using SYBR Green PCR Core Reagents (Applied Biosystems). Reaction mixtures were incubated for 10 min at 95 °C, followed by 40 cycles of 15 s at 95 °C, 1 min at 60 °C, and finally 15 s at 95 °C, 1 min 60 °C, and 15 s at 95 °C. For each mRNA, gene expression was normalized to the ribosomal protein S11 (rps11) content in each sample using the Pfaffl method ^34^. The primers used are shown in **Table S1**. In all cases, each PCR was performed with triplicate samples and repeated at least with two independent samples.

### Chemical treatments

In some experiments, 24-hours post fertilization (hpf) embryos manually dechorionated were treated with caspase-3 inhibitor (C3I, 50 µM, MCE), plasmalogens or ether lipid precursor (20 µM, Avanti lipids) from 1-5 days by bath immersion at 28 °C. Incubation was carried out in 6-well plates containing 20 to 25 larvae/well in egg water supplemented with 1% dimethyl sulfoxide (DMSO) as vehicle. Plasmalogens and the custom-synthesized ether lipid precursor were dried under a N_2_ stream at 35 °C and resuspended in DMSO: HsVEPE1 (1-(1*Z*-octadecenyl)-2-oleoyl-*sn*-glycero-3-phosphoethanolamine or 18(Plasm)-18:1 PE), HsVEPE2 (1-(1*Z*-octadecenyl)-2-arachidonoyl-*sn*-glycero-3-phosphoethanolamine or 18(Plasm)-20:4 PE), HsVEPE3 (1- (1*Z*-octadecenyl)-2-docosahexaenoyl-*sn*-glycero-3-phosphoethanolamine or 18(Plasm)-22:6 PE), HsAEPE1 (1-*O*-octadecanyl-2-oleoyl-*sn*-glycero-3-phosphoethanolamine or *O*-18:0-18:1 PE).

### In vivo imaging of zebrafish larvae

To study the total number, the distribution and recruitment of immune cells, 2 or 3-days post fertilization (dpf) transgenic larvae were anesthetized in embryonic medium with 0.16 mg/ml tricaine. Images were taken of the areas of interest (otic region and tail) or of the whole body at different time points, as appropriate considering the study model, using a Leica MZ16F fluorescence stereomicroscope. The number of fluorescent cells was determined by visual counting, and the fluorescence intensity of different parameters (*il1b*, *nfkb*, and H_2_O_2_) was obtained and analyzed from a region of interest (ROI) with ImageJ (FIJI) software ^35^.

### Tail transection and regeneration assay

For tail transection experiments, 3 dpf larvae anesthetized by tricaine bath immersion were transected from the tail with a sterile micro-scalpel as previously described ^36^. At least three independent experiments were performed with a total number of 20-50 larvae per treatment. In vivo neutrophil and macrophage recruitment was evaluated by assessing the recruitment of cells at wound sites (the region posterior to the circulatory loop) at various time points throughout inflammatory process. Macrophages polarization was analyzed by the doble positive fluorescent signal using *Tg(mfap4.1:mCherry-F)* and *Tg(tnfa:eGFP)* transgenic lines. Tissue regeneration was also evaluated by assessing the regenerated tail fin area at 3, 6, 24 and 48-hours post-wounding (hpw) using Image J software. Regeneration was calculated by analyzing the regenerated tail fin area vs. fin areas of unamputated animal.

### Oxidative stress assay

H_2_O_2_ release at the wound site was quantified employing the live cell fluorogenic substrate acetyl-pentafluor-obenzene sulphonyl fluorescein (Cayman Chemical, Ann Arbor, Michigan, USA), as previously described ^37^. Briefly, larvae of 3 dpf were incubated with 50 µM of the substrate reagent prior tail fin amputation procedure for 1 hour in a 24-well plate, then immediately injured. To determine oxidative response post injury at the wound intensity of fluorescence was assessed using Image J software.

### Infection assay

For the infection experiments, *Salmonella enterica* serovar Typhimurium strain 12023 (wild type, WT) was used. Overnight cultures in Luria-Bertani (LB) broth were diluted 1/5 in LB with 0.3 M NaCl, incubated at 37 °C until 1.5 optical density at 600 nm was reached, and finally diluted in sterile PBS. Larvae of 2 dpf were anaesthetized in embryo medium with 0.16 mg ml−1 tricaine and 10 bacteria (yolk sac, systemic infection) or 200 (otic vesicle, myeloid cell recruitment) per larvae were microinjected. Larvae were allowed to recover in egg water at 28–29 °C and monitored for clinical signs of disease or mortality over 5 days. Neutrophil and macrophage recruitment to the otic vesicle was analyzed as described above for tail wounding. At least three independent experiments were performed with a total number of 20-100 larvae per treatment.

### Statistical analysis

Statistical analysis was performed using Prism 8.0 (GraphPad Software, CA, USA). Data are shown as mean ± s.e.m. and were analyzed by analysis of variance and a Tukey multiple range test to determine differences between groups. The differences between two samples were analyzed by the Student’s *t*-test. A log-rank test was used to calculate the statistical differences in the survival of the different experimental groups.

### Data sharing statement

For original data, please contact.

## Results

### Inhibition of plasmalogen synthesis promotes a proinflammatory stage with increased neutrophil and macrophage apoptosis

To characterize the specific role of plasmalogens *in vivo* under physiological conditions, single-cell stage eggs were microinjected with specific crRNA/Cas9 complexes (cr*peds1a* and cr*peds1b*) to inhibit the two zebrafish *peds1* paralogs. High *peds1* knockdown efficiency was achieved with a concomitant reduction of total plasmalogen levels in cr*peds1* larvae (estimated by measuring the dimethylacetal or DMA derivatives; see Materials and Methods), which was more pronounced at 5 dpf (Supplementary Figure 1). This may be related to maternal transfer of plasmalogens, since we found that plasmalogen content in control larvae was 2-fold higher at 5 dpf than at 3 dpf, suggesting that their biosynthesis begins between 3 and 5 dpf. Specific DMAs were either decreased (C16:0, C17:0, C:18:0) or became undetectable (C18:1, C20:0) in Peds1-deficient larvae relative to control larvae (Supplementary Figure 1B).

Peds1-deficient larvae were slightly smaller than control larvae at both 3 and 5 dpf (Figure 1A), but they did not show any obvious developmental defect. Notably, Peds1 deficiency resulted in robust induction of the inflammation reporters Nfkb and Il1b (Figure 1B and C). In agreement, increased transcript levels of *nfkb1* and a strong induction of *il1b, tnfa* and *cxcl8a* expression was also observed when plasmalogen synthesis was inhibited (Figure 1D). Moreover, and surprisingly, the total number of neutrophils and macrophages was lower in cr*peds1* larvae than in their wild-type siblings, and the observed neutropenia and monocytopenia was fully reversed upon pharmacological inhibition of caspase-3 (Figure 1E-F). However, there were no differences between the two larval groups in the numbers of hematopoietic stem and progenitor cells (HSPCs) (Figure 1G) or of erythrocytes (Figure 1H).

**Figure 1.**
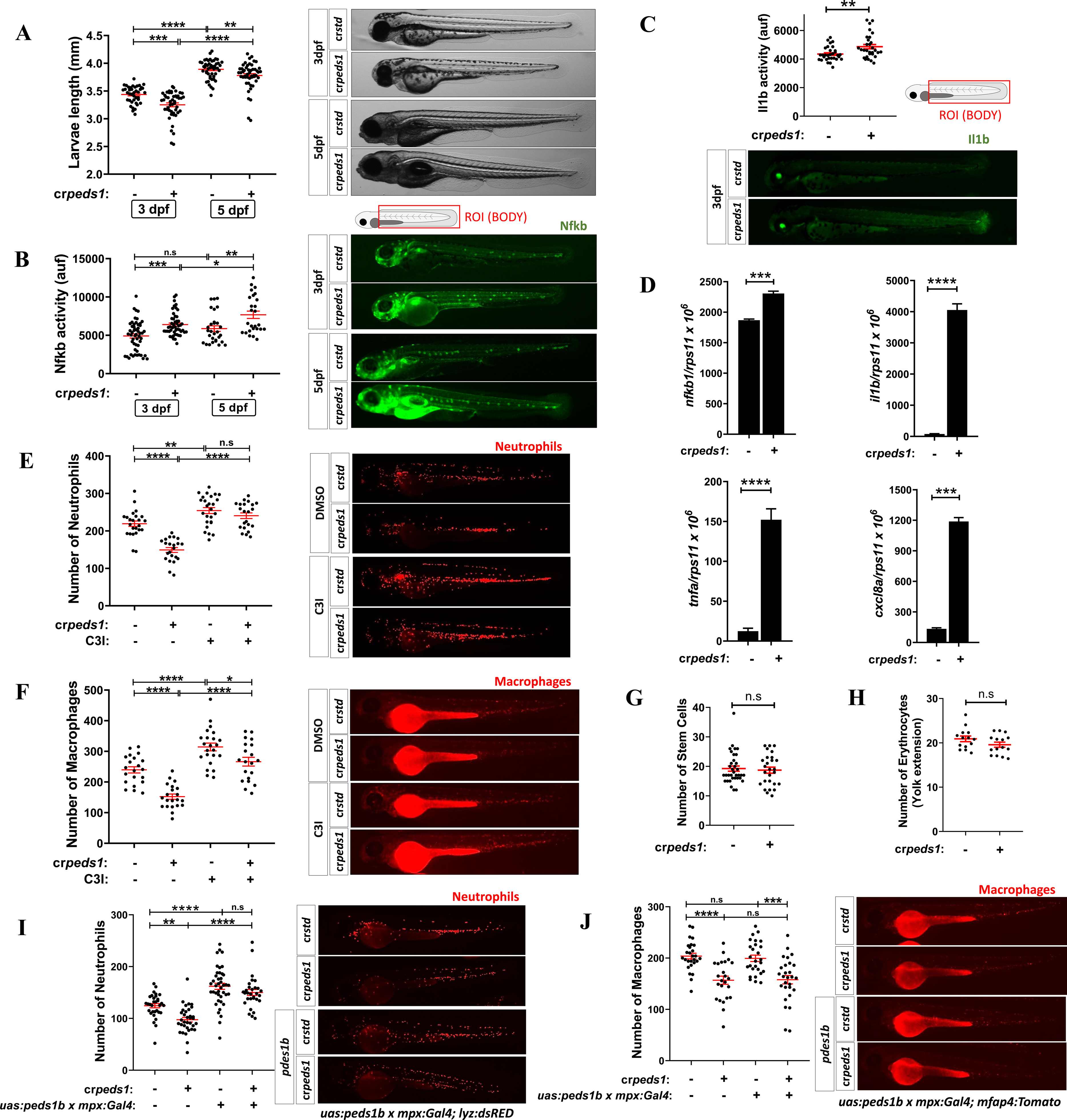
Peds1 deficiency promotes a proinflammatory stage with increased neutrophil and macrophage apoptosis. One-cell stage zebrafish eggs were microinjected with *std* or *peds1* crRNA/Cas9 complexes and the following parameters were evaluated: (A) Larval length and representative images (brightfield channel) at 3- and 5 dpf, (B) Nfkb activity and representative images (green channel; *Tg(NFkB-RE:eGFP)*) at 3 and 5 dpf analyzed by fluorescence microscopy, (C) Il1b activity and representative images (green channel; *Tg(il1b:GFP-F)^ump3^*) at 3 dpf analyzed by fluorescence microscopy. (D) Transcript levels of the indicated genes analyzed at 3 dpf by RT-qPCR. (E-F) Number of neutrophils and macrophages and representative images of microinjected larvae after 48 hour of treatment with caspase 3 inhibitor (C3I) or vehicle (DMSO) were quantitated by fluorescence microscopy (red channel; *Tg(lyz:DsRED2)* and *Tg(mfap4.1:mCherry-F)*, respectively). (G-H) Hematopoietic stem and progenitor cells (green channel; *Tg(-6.0itga2b:eGFP)*) and erythrocytes (red channel: *Tg(gata1a:dsRED)*) number were quantitated by fluorescence microscopy. (I-J) Number of neutrophils and macrophages and representative images of larvae microinjected to force expression of *peds1* in neutrophils using the neutrophil-specific myeloperoxidase (*mpx*) promoter and analyzed by fluorescence microscopy. Each point represents one larva and the mean ± SEM of each group is also shown. *P* values were calculated using one-way ANOVA Tukey’s and multiple range or unpaired Student’s *t*-test. n.s, not significant, * *p* ≤ 0.05, \*\**p* ≤ 0.01, \*\*\**p* ≤ 0.001, \*\*\*\**p* ≤ 0.001. auf: arbitrary units of fluorescence.

We next wondered whether the effect of Peds1 deficiency in neutrophils and macrophages was cell autonomous. The results showed that forced expression of *peds1* in neutrophils using the neutrophil-specific myeloperoxidase (*mpx*) promoter was able to restore the number of neutrophils (Figure 1I), but not of macrophages (Figure 1J), in Peds1-deficient larvae. Collectively, these results suggest a cell autonomous effect of Peds1 in myeloid cells and the non-transferability of plasmalogens between these two cell types.

### Inhibition of plasmalogen synthesis exacerbates chronic skin inflammation

The above results prompted us to study the impact of plasmalogens in chronic inflammation using Spint1a-deficient larvae, which are characterized by the presence of skin lesions or keratinocyte aggregates, neutrophil mobilization from the caudal hematopoietic tissue (CHT) to inflamed skin, and neutrophilia (Figure 2A) ^38,39^. We observed again that Peds1-deficient larvae were slightly smaller than their control counterparts (Figure 2B). In addition, Peds1 deficiency led to increased skin lesions (Figure 2C) and Nfkb activity (Figure 2D) at 3 dpf and, even at 5 dpf, when the skin inflammation was largely resolved in control larvae (Supplementary Figures 2A, 2B). Moreover, Peds1-deficient larvae had increased neutrophil infiltration into the skin both at 3 and 5 dpf (Figure 2E and Supplementary Figure 2C), even though they had lower number of neutrophils than control larvae (Figure 2E).

**Figure 2.**
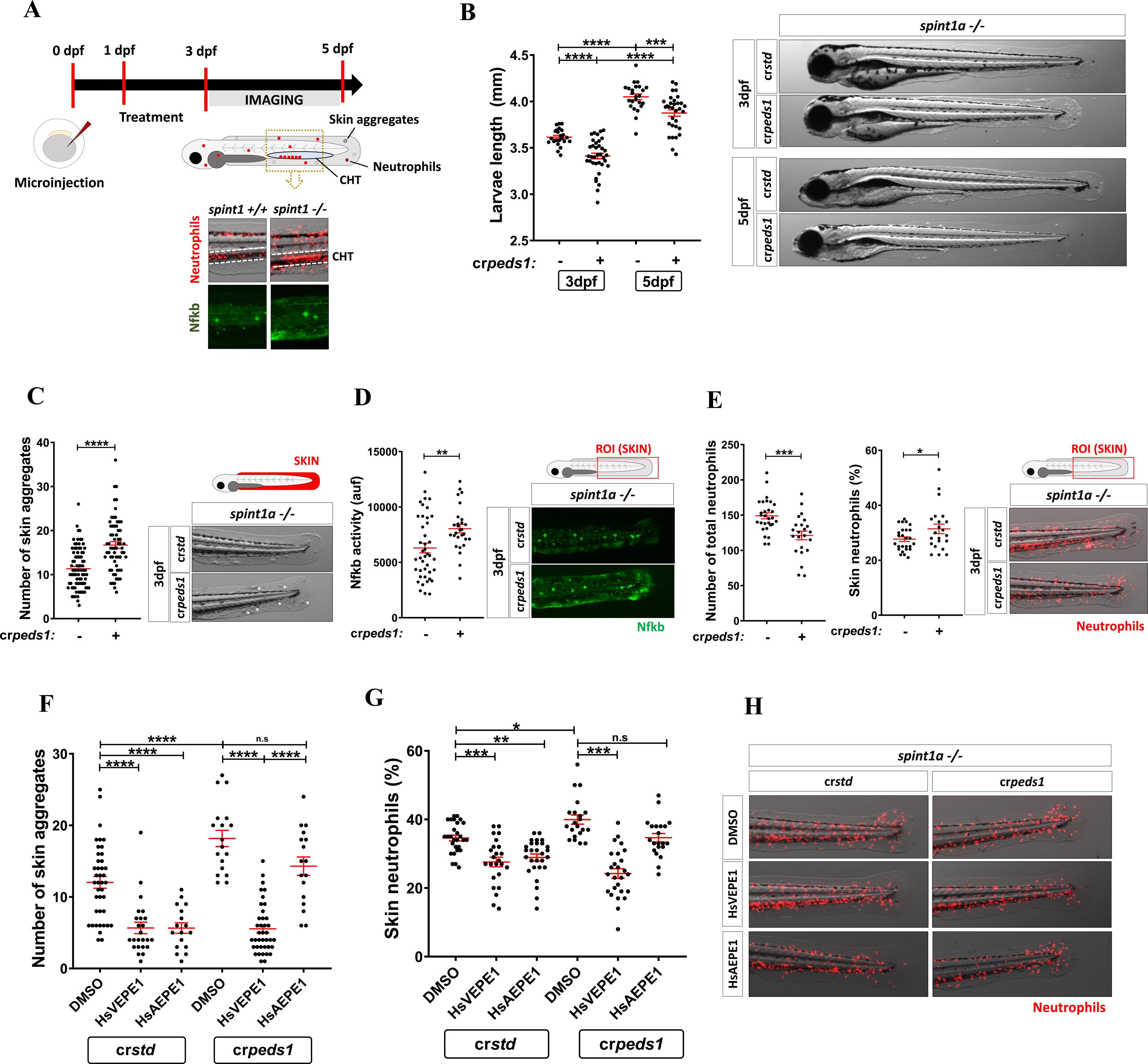
Peds1 deficiency exacerbates skin chronic inflammation. (A) Schematic of the experimental procedure used for skin chronic inflammation assays. Spint1a-deficient one-cell stage eggs were microinjected with *std* or *peds1* crRNA/Cas9 complexes and imaged at 3 and 5 dpf. Treatments of interest were added to dechorionated embryos at 1 dpf by bath immersion and renewed daily. Spint1a-deficient larvae showed chronic skin inflammation with increased skin neutrophil infiltration, Nfkb activity and keratinocyte aggregates. (B) Analysis and representative images (brightfield channel) of larval length at 3 and 5 dpf. (C) Number of keratinocyte aggregates in the skin (white arrows) and representative images (brightfield channel) at 3 dpf. (D) Analysis and representative images of Nfkb activity (green channel; *Tg(NFkB-RE:eGFP)^sh235^*) at 3 dpf by fluorescence microscopy. (E) Total number and percentage of neutrophils in the skin and representative merge images (brightfield and red channel; *Tg(lyz:DsRED2)^nz50^*) quantitated by fluorescence microscopy. (F) Number of skin aggregates and (G) percentage of skin neutrophils quantitated in cr*std* and cr*peds*1 *spint1a-/-* larvae treated for 48 hours with 20 µM HsVEPE1, 20 µM HsAEPE1 or vehicle (DMSO). (H) Representative merge images (brightfield and red channel; *Tg(lyz:DsRED2)^nz50^*) of each treatment analyzed in F and G by microscopy fluorescence are shown. Each point represents one larva and the mean ± SEM of each group is also shown. *P* values were calculated using one-way ANOVA and Tukey’s multiple range or unpaired Student’s *t*-test, as appropriate. n.s, not significant, * *p* ≤ 0.05, \*\**p* ≤ 0.01, \*\*\**p* ≤ 0.001, \*\*\*\**p* ≤ 0.001. auf: arbitrary units of fluorescence.

Since inhibition of plasmalogen biosynthesis worsened skin inflammation in Spint1a-deficient larvae, we tested the effect of exogenous supplementation of dechorionated embryos with a commercially available human plasmalogen (HsVEPE1) that has a single double bond at the *sn*-2 position (see Materials and Methods). First, we confirmed that Spint1a-deficient larvae bathed in 20 µM of HsVEPE1 were capable of incorporating the plasmalogen, as judged by the increased C18:0 DMA levels observed (Supplementary Figure 2D). Importantly, this concentration of HsVEPE1 alleviated skin inflammation, assayed as the number of keratinocyte aggregates and neutrophil infiltration (Supplementary Figure 2E). Subsequently, we performed the same experiment but adding HsVEPE1 or its alkyl ether precursor HsAEPE1 to Peds1-deficient and control larvae. Interestingly, both HsVEPE1 and HsAEPE1 decreased skin lesions and neutrophil infiltration in control larvae (Figure 2F-2H). However, in Peds1-deficient larvae, only HsVEPE1 was able to decrease keratinocyte aggregates and neutrophil infiltration (Figure 2F-2H) and to rescue larval length (Supplementary Figure 2F), likely due to the inability of these larvae to convert HsAEPE1 into HsVEPE1. As the presence of PUFAs in plasmalogens may increase their antioxidant properties, we also tested two plasmalogens enriched in PUFAs, namely HsVEPE2 and HsVEPE3 (with four and six double bonds, respectively, in the acyl chain at *sn*-2). Both plasmalogens ameliorated skin inflammation in Spint1a-deficient larvae, but to levels that resembled those observed with HsVEPE1 (Supplementary Figure 2G). Overall, these results demonstrate the anti-inflammatory function of plasmalogens, but not of their precursors, in this chronic skin inflammation model.

### Inhibition of plasmalogen synthesis impairs inflammation resolution and tissue regeneration

The effect of inhibiting plasmalogen synthesis under acute sterile inflammation was also evaluated. For this purpose, Peds1-deficient larvae were tail-cut and neutrophil and macrophage recruitment, oxidative stress and inflammation were analyzed (Figure 3A). Although recruitment to wound at early times (3- and 6-hours post-wounding, hpw) was normal in Peds1-deficient larvae, at 24 and 48 hpw they showed higher neutrophil (Figure 3B) and lower macrophage (Figure 3C) recruitment than control larvae. However, the number of proinflammatory macrophages, i.e. M1-like that express Tnfa ^28^, was higher in Peds1-deficient larvae at all times evaluated (Figure 3D). Oxidative stress, as assessed by hydrogen peroxide release, was found to be higher in larvae deficient in Peds1 at 3 hpw. In agreement with the above data, Peds1 deficiency caused a dramatic increase in Il1b (Figure 3F) and Nfkb (Figure 3G) activity at 24 and 48 hpw. These results suggest that Peds1-deficient larvae fail to resolve inflammation.

**Figure 3.**
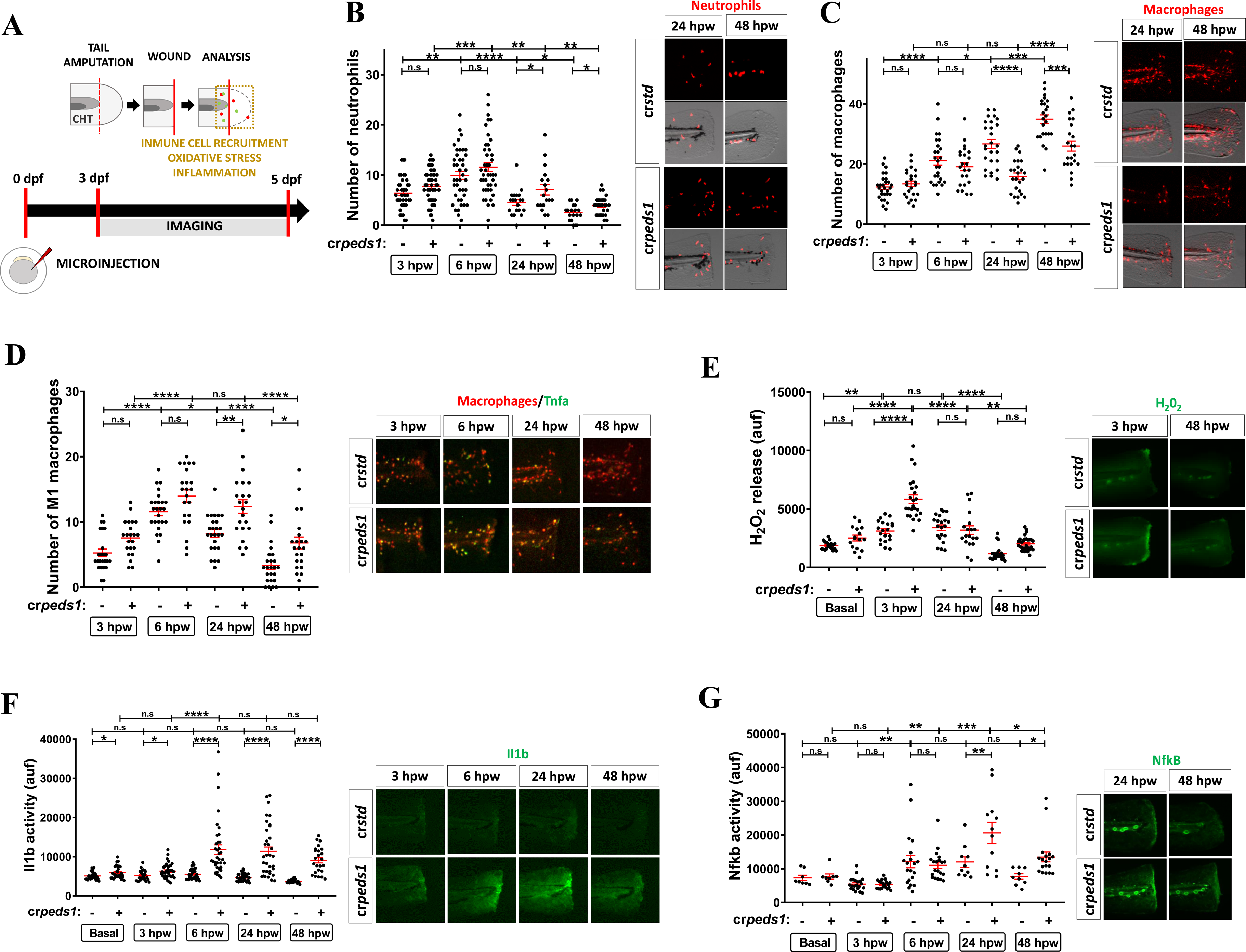
Peds1 deficiency impairs inflammation resolution and causes aberrant immune cells recruitment in a sterile tail injury model. (A) Schematic of the experimental procedure used for tail injury assays. One-cell stage zebrafish eggs were microinjected with *std* or *peds1* crRNA/Cas9 complexes. At 3dpf larvae were tail amputated and then inflammation and immune cell recruitment at the injury site (dotted lines) were analyzed at 3, 6, 24 and 48 hours post-wounding (hpw) by fluorescence microscopy. The wound area is defined as the region between the amputation edge and the end of the caudal hematopoietic tissue (CHT). (B-C) Number of neutrophil and macrophages recruited at the injury site and representative merge images (brightfield and red channel; *Tg(lyz:DsRED2)^nz50^* and *Tg(mfap4.1:mCherry-F)^xt12^*, respectively). (D) Number of M1-like macrophages (Tnfa^+^) recruited at the injury site and representative merge images (red channel: *Tg(mfap4.1:mCherry-F)^xt12^*and green channel: *Tg(tnfa:eGFP-F)^ump5^*). (E) Analysis of H_2_O_2_ release at the wound site using the fluorogenic substrate acetyl-pentafluor-obenzene sulphonyl fluorescein. (E-G) Analysis and representative images of Il1b activity (green channel; *Tg(il1b:GFP-F)^ump3^*) and Nfkb activity (green channel; *Tg(NFkB-RE:eGFP)^sh235^*) at the wound site. Each point represents one larva and the mean ± SEM of each group is also shown. *P* values were calculated using one-way ANOVA and Tukey’s multiple range. n.s, not significant, * *p* ≤ 0.05, \*\**p* ≤ 0.01, \*\*\**p* ≤ 0.001, \*\*\*\**p* ≤ 0.001. auf, arbitrary units of fluorescence.

Since Peds1 deficiency altered the inflammatory response to induced damage, its effect on tissue regeneration was next determined (Figure 4A). Unlike control larvae, which completely regenerated the tail at 48 hpw, Peds1-deficient larvae failed to do so (Figure 4B). Therefore, we wondered whether this impaired regeneration of Peds1-deficient larvae was due to failure in resolving inflammation or, alternatively, to plasmalogens being required for cell proliferation. As Il1b was dramatically induced in the wound of Peds1-deficient larvae, but it hardly increased in control larvae, we analyzed the effect of knocking down this proinflammatory cytokine (knockout efficiency of about 80%) (Supplementary Figure 3). While Il1b deficiency had no effect on tail regeneration of control larvae, it fully rescued regeneration and inflammation, assayed as Il1b reporter activity, in Peds1-deficient larvae (Figure 4C, 4D). Therefore, the inability of Peds1-deficient larvae to resolve inflammation appears to be responsible for the impaired tissue regeneration in this acute inflammation model.

**Figure 4.**
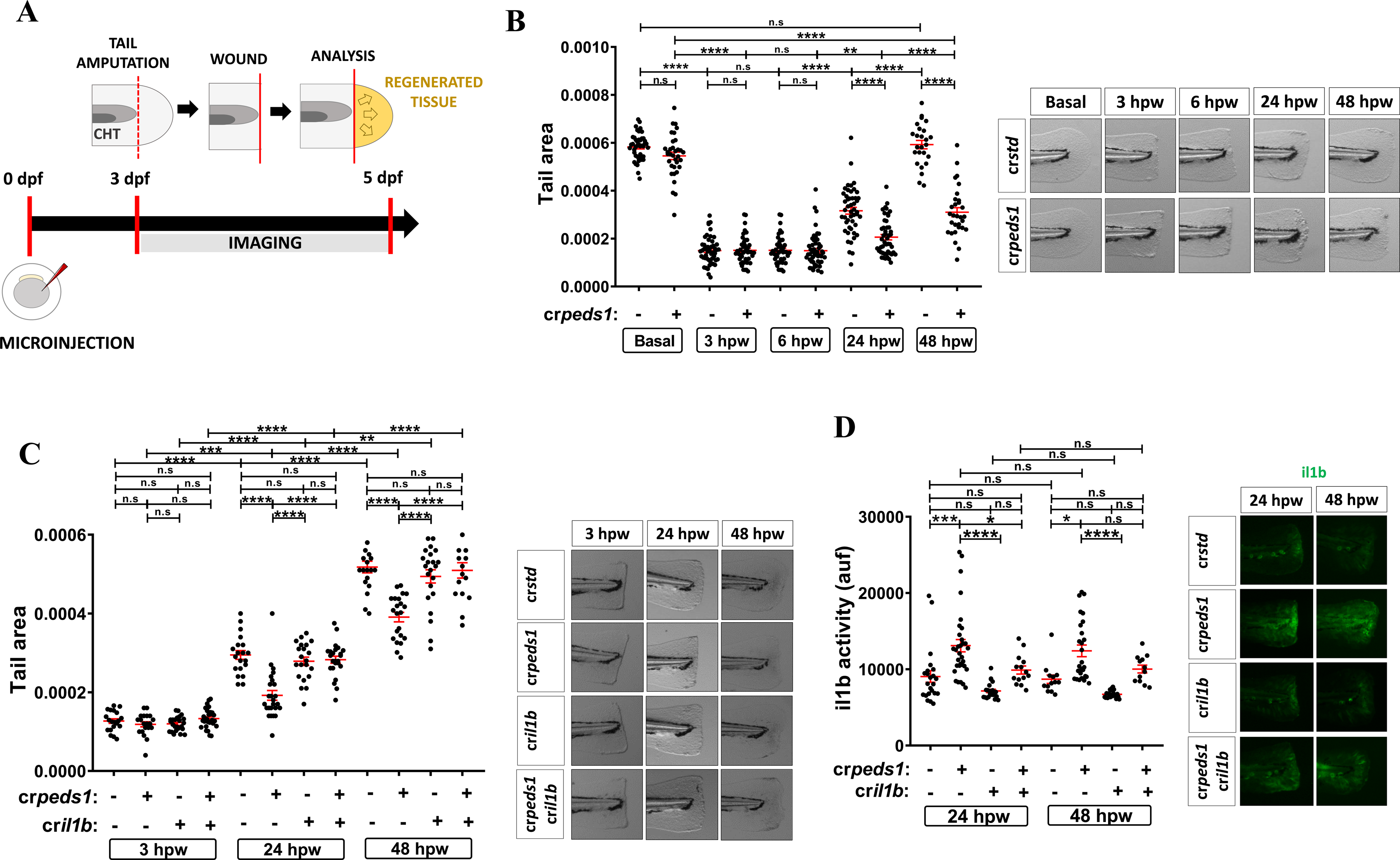
Peds1 deficiency hampers tissue regeneration. (A) Schematic of the experimental procedure used for tail injury and regeneration assays. Three dpf larvae previously microinjected with *std* or *peds1* crRNA/Cas9 complexes alone or in combination with cr*il1b* were tail fin amputated and regeneration was evaluated by measuring regenerated fin areas up to 5 dpf (yellow). (B-C) Measurement of regenerated fin areas and representative images (brightfield channel). (D) Analysis and representative images of Il1b activity (green channel; *Tg(il1b:GFP-F)^ump3^*) by microscopy fluorescence. Each point represents one larva and the mean ± SEM of each group is also shown. *P* values were calculated using one-way ANOVA and Tukey’s multiple range. n.s, not significant, * *p* ≤ 0.05, \*\**p* ≤ 0.01, \*\*\**p* ≤ 0.001, \*\*\*\**p* ≤ 0.001. auf, arbitrary units of fluorescence.

### Inhibition of plasmalogen synthesis results in increased susceptibility to infection

The effect of inhibiting plasmalogen synthesis on the resistance to infection by the intracellular bacterium *Salmonella enterica* serovar Thyphimurium (ST), upon microinjecting it in yolk sac, was also evaluated (Figure 5A). The results showed that Peds1-deficient larvae are more susceptible to ST than their wild type siblings (Figure 5B). Notably, supplementation with exogenous plasmalogen (HsVEPE1) not only mitigated the higher susceptibility of Peds1-deficient larvae to ST infection, but it also robustly increased the resistance of wild type larvae (Figure 5B). As expected, supplementation with the plasmalogen precursor (HsAEPE1) increased infection resistance of control larvae but failed to alleviate the higher susceptibility to ST of Peds1-deficient larvae (Figure 5C). Since neutrophils play a critical role in the resistance of zebrafish larvae to ST ^40^, we tested the effects of inhibiting their apoptosis in infected Peds1-deficient larvae and found that the caspase-3 inhibitor phenocopied the effects of plasmalogen supplementation in larval survival (Figure 5D). Furthermore, inhibition of caspase-3 also rescued neutrophil recruitment to the otic vesicle of infected Peds1-deficient larvae (Figure 5E). As regards, the impact of neutrophil apoptosis inhibition in total neutrophil number of Peds1-deficient larvae, Peds1 deficiency impaired the early neutrophilia induced by ST infection and caspase-3 inhibition was able to normalize the neutrophil count (1 hpi, Figure 5F). Finally, the relevance of neutropenia in the higher susceptibility of Peds1-deficient larvae to ST was further confirmed by the ability of forced expression of granulocyte colony-stimulating factor (*csf3a*), which induces neutrophil production, to robustly increase the resistance of wild type larvae and partially rescue the resistance (Figure 5G) and neutrophil number of Peds1-deficient larvae (Figure 5H).

**Figure 5.**
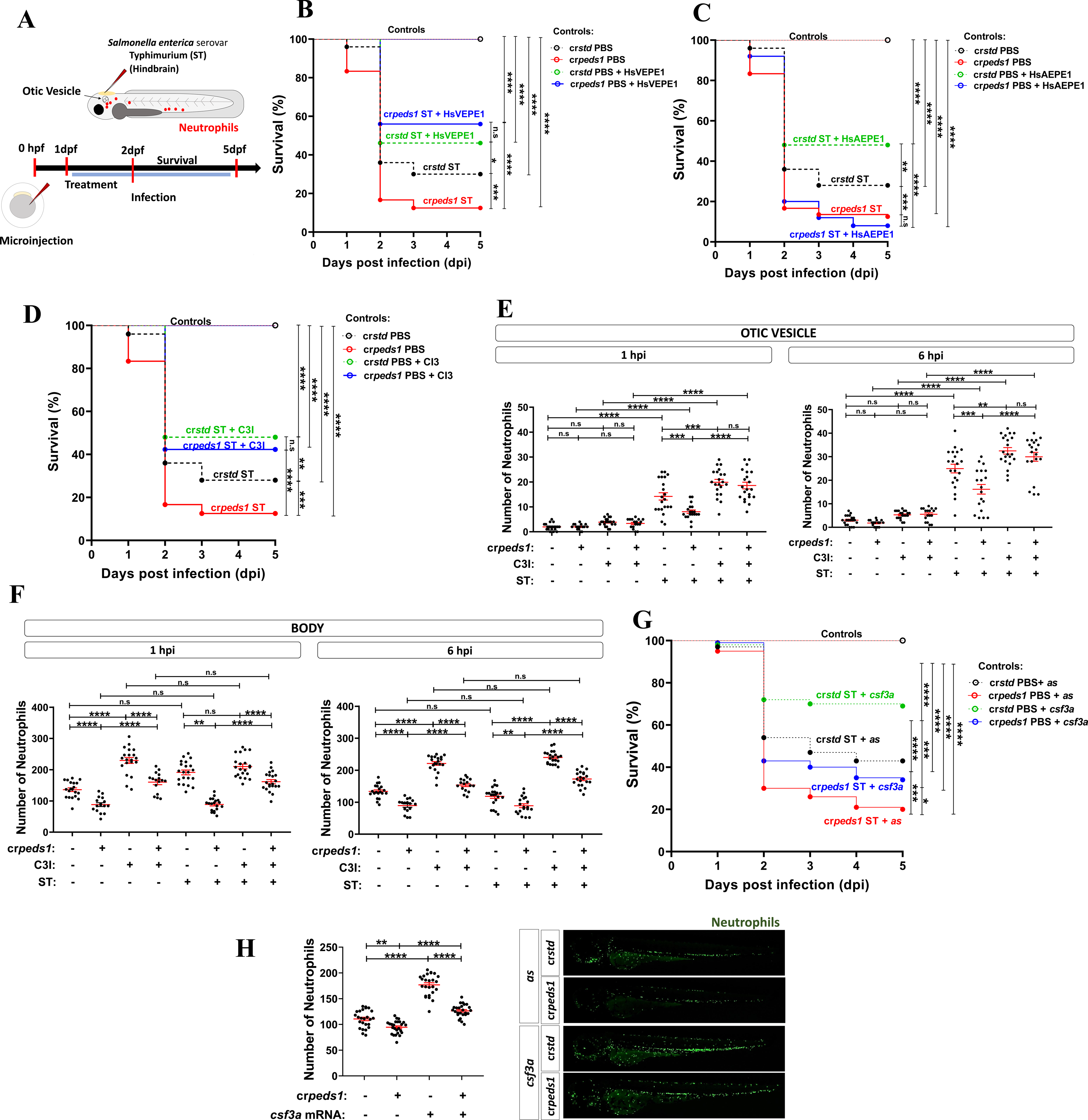
Peds1 deficiency increased susceptibility to bacterial infection. (A) Schematic of the experimental procedure used for infection assays. Single-cell zebrafish embryos microinjected with *std* or *peds1* crRNA/Cas9 complexes were dechorionated and infected at 2 dpf through the hindbrain with *Salmonella enterica* serovar Thyphimurium (ST). Treatments of interest were added daily from 1dpf by immersion and the number of surviving larvae was counted daily for the next 5 days. (B) Survival analysis of cr*std* and cr*peds1* larvae infected with ST and treated with 20 µM HsVEPE1 or vehicle (DMSO). (C) Survival analysis of cr*std* and cr*peds1* larvae infected with ST and treated with 20 µM HsAEPE1 or vehicle (DMSO). (D) Survival analysis of cr*std* and cr*peds1* larvae infected with ST and treated with caspase 3 inhibitor (C3I, 50 µM) or vehicle (DMSO). (E-F) Number of neutrophil in the otic vesicle and body at 1 and 6 hour post-infection in cr*std* and cr*peds1* larvae treated with 50 µM C3I or vehicle (DMSO) counted by fluorescence microscopy (red channel; *Tg(lyz:DsRED2)^nz50^*). (G) Survival analysis of cr*std* and cr*peds1* larvae injected in combination with antisense (As) or *csf3a* mRNA and infected with ST. (H) Number of neutrophil in whole body in uninfected cr*std* and cr*peds1* larvae of 3 dpf treated with 50 µM C3I or vehicle (DMSO) counted by fluorescence microscopy (green channel; *(mpx:eGFP)^i114^*). (B-D and G) A log-rank test was used to calculate the statistical differences in the survival of the different experimental groups. (E-F) Each point represents one larva and the mean ± SEM of each group is also shown. *P* values were calculated using one-way ANOVA and Tukey’s multiple range. n.s, not significant, * *p* ≤ 0.05, \*\**p* ≤ 0.01, \*\*\**p* ≤ 0.001, \*\*\*\**p* ≤ 0.001.

## Discussion

The role of plasmalogens in inflammation remains controversial, largely because until recently most studies could not distinguish the individual contributions of plasmalogens from those of their alkyl ether lipids precursors. The present study reveals the relevance of endogenous plasmalogens in vertebrate myeloid cell biology and inflammation (Figure 6). Peds1 deficiency resulted in exacerbated inflammation and delayed development in zebrafish larvae, without provoking apparent morphological alterations. While HSPC emergence and erythropoiesis proceeded normally in Peds1-deficient larvae, they suffered from robust neutropenia and monocytopenia that was rescued by pharmacological inhibition of caspase-3. Forced expression of *peds1b* in neutrophils restored neutrophil, but not macrophage, number in Peds1-deficient larvae, suggesting a cell autonomous effect of Peds1 in myeloid cells and the non-transferability of plasmalogens from neutrophils to macrophages. It has been previously reported that bone marrow-specific deletion of either fatty acid synthase (FAS), which catalyzes the first committed step in de novo lipogenesis, or of peroxisomal reductase activating PPARγ (PexRAP), which acts in ether lipid synthesis following generation of the ether bond by AGPS at the peroxisome, result in neutropenia, but not monocytopenia, due to cell-autonomous loss of neutrophils via endoplasmic reticulum stress and apoptosis ^41^. However, since neutropenia was not observed in other mouse models of ether lipid deficiency, such as GNPAT knockout mice, or in patients suffering from RCDP, it was argued that accumulation of the intermediate 1-O-alkyl-glycerone phosphate or tamoxifen toxicity might be responsible for neutrophil death observed in PexRAP-deficient mice ^42^. Our results unequivocally show that endogenous plasmalogen biosynthesis is required to maintain neutrophil and macrophage viability in zebrafish larvae.

**Figure 6.**
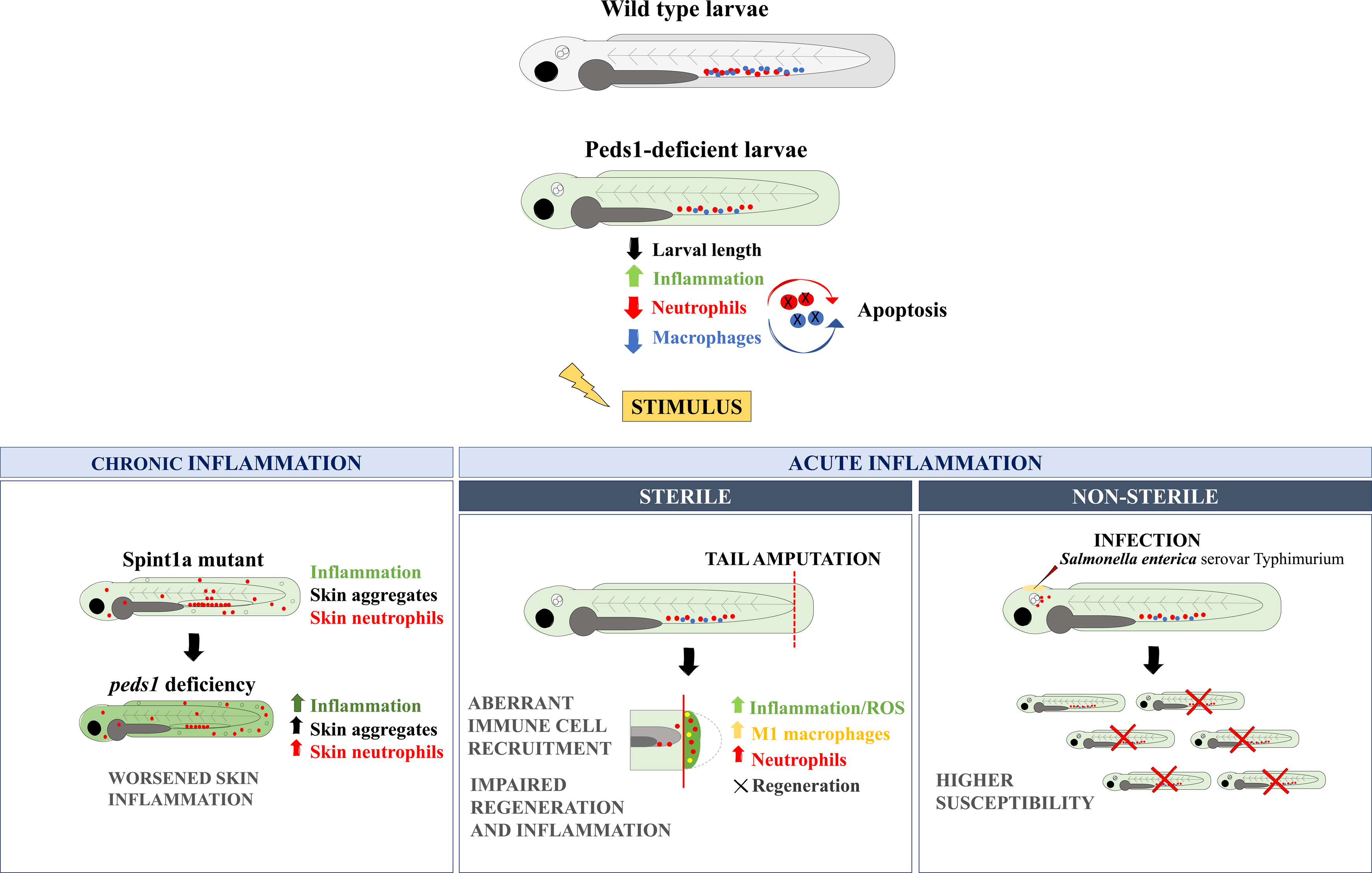
Model showing the effect of inhibiting plasmalogen synthesis under basal and different inflammatory conditions. Inhibition of plasmalogen synthesis promotes a proinflammatory stage with increased neutrophil and macrophage apoptosis. Plasmalogen deficiency also exacerbates chronic skin inflammation, impairs inflammation resolution and tissue regeneration, and results in increased susceptibility to infection.

Importantly, our study reveals that Peds1 deficiency in zebrafish exacerbates both acute and chronic inflammation (Figure 6), hinting that the diminished plasmalogen levels observed in chronic inflammation-linked diseases might be a cause rather than a consequence of these diseases. Moreover, exogenous addition of plasmalogens not only rescued the exacerbated inflammation of Peds1-deficient larvae, but also reduced inflammation in wild type animals, thus providing additional support for the anti-inflammatory role exerted by PRT in several animal disease models and patient-derived cells studies, and in AD and PD patients (Bozelli et al., 2021; Dorninger et al., 2020). Our results may also have clinical relevance in PRT, since plasmalogens, but not their precursors, mediate the anti-inflammatory effects in zebrafish, as revealed by the failure of the precursors to alleviate inflammation in Peds1-deficient larvae. Our finding that all the plasmalogens used, regardless of the number of double bonds in their acyl chain at *sn*-2, similarly ameliorated skin inflammation, rules out that the observed effects were related to their PUFA content.

The acute inflammation model revealed a critical role of endogenous plasmalogens in myeloid cell activation. Although Peds1-deficient larvae were neutropenic and monocytopenic, early neutrophil and macrophage recruitment to wound was normal. However, inflammation resolution was impaired, as evidenced by neutrophil and Tnfa^+^ macrophage persistence in the wound, and dramatic increased and lasting levels of H_2_O_2_, Il1b and Nfkb in the wound area (Figure 6). Strikingly, we observed impaired tissue regeneration in Peds1-deficient larvae, which was fully rescued by Il1b deficiency. The role played by Il1b in tail regeneration is complex. While a transient production of Il1b by epithelial cells appears to be required for proper regeneration through the induction of genes encoding important regenerative factors, exacerbated production of Il1b promotes the apoptosis of regenerative cells ^43,44^. In addition, macrophages are required for the attenuation of Il1b production in the wound, thus preventing apoptosis of regenerative cells ^43^. Furthermore, the reduced number of recruited macrophages at later stages of regeneration and their skew towards M1-like phenotype may further impair the regeneration process ^45^. Whatever the outcome, our results point to a critical role for plasmalogens in tissue regeneration by promoting the resolution of inflammation rather than by a direct effect on cell proliferation and differentiation.

Our results also revealed the importance of plasmalogens in the response to bacterial infection (Figure 6). Hence, Peds1-deficient larvae were more susceptible to bacterial infection but, more importantly, exogenous plasmalogens not only alleviated this higher susceptibility of Peds1-deficient larvae, but also increased the resistance of wild-type larvae. These effects were again mediated by plasmalogens, since their precursor increased bacterial resistance of wild-type but not of Peds1-deficient larvae. Furthermore, the higher susceptibility of Peds1-deficient larvae to bacterial infection was fully rescued by pharmacological inhibition of caspase-3 and partially rescued by Csf3a-induced myelopoiesis. These results further suggest that plasmalogens are required for neutrophil survival rather than migration, activation and microbicidal functions. The role of plasmalogens in human myeloid cell biology is largely unknown. It has been reported that neutrophil activation results in changes in plasma membrane phospholipids, including phosphatidylethalonamine plasmalogen content ^46^. Similarly, the plasmalogen profile was also altered during the differentiation of monocytes to macrophages ^47^. In addition, human neutrophils include a unique plasmenylethalonamine pool that undergoes myeloperoxidase-dependent oxidation during activation and is elevated in the plasma during sepsis ^48^.

In summary, we report for the first time a cell autonomous role of plasmalogens in the survival of vertebrate neutrophils and macrophages, which seems to be dispensable for their recruitment and activation. In addition, while endogenous plasmalogen production was found to be critical to regulate inflammation and promote its resolution, exogenous plasmalogen supplementation exerted beneficial effects by dampening physiological and pathological inflammation and increasing bacterial infection clearance. The zebrafish model developed here is a unique tool to shed light into the relevance of plasmalogens in physiology and disease.

## Acknowledgments

We thank I. Fuentes and P.J. Martínez for their excellent technical assistance, and S.A. Renshaw, P. Crosier, D. Tobin, and L.I. Zon for the zebrafish lines, and D. Holden for the bacterial strain.

## Authorship and conflict-of-interest statements

ABA, DGM, MEA and VM conceived the study; ABA, SDT, EBM, AJMG, JCS and MBC performed the research; ABA, SDT, EBM, AJMG, JCS, MBC, DGM, MEA and VM analyzed data; ABA, EBM and VM wrote the original draft; MEA and VM edited the final version. All authors have read and agreed to the published version of the manuscript.

The authors declare no conflict-of-interest.

## References

1. Harayama T, Riezman H. Understanding the diversity of membrane lipid composition. Nat Rev Mol Cell Biol. 2018;19(5):281–296.

2. Jimenez-Rojo N, Riezman H. On the road to unraveling the molecular functions of ether lipids. FEBS Lett. 2019;593(17):2378–2389.

3. Padmanabhan S, Monera-Girona AJ, Pajares-Martinez E, et al. Plasmalogens and Photooxidative Stress Signaling in Myxobacteria, and How it Unmasked CarF/TMEM189 as the Delta1’-Desaturase PEDS1 for Human Plasmalogen Biosynthesis. Front Cell Dev Biol. 2022;10:884689.

4. Wainberg M, Kamber RA, Balsubramani A, et al. A genome-wide atlas of co-essential modules assigns function to uncharacterized genes. Nat Genet. 2021;53(5):638–649.

5. Werner ER, Keller MA, Sailer S, et al. The TMEM189 gene encodes plasmanylethanolamine desaturase which introduces the characteristic vinyl ether double bond into plasmalogens. Proc Natl Acad Sci U S A. 2020;117(14):7792–7798.

6. Gallego-Garcia A, Monera-Girona AJ, Pajares-Martinez E, et al. A bacterial light response reveals an orphan desaturase for human plasmalogen synthesis. Science. 2019;366(6461):128-132.

7. Bozelli JC, Jr., Azher S, Epand RM. Plasmalogens and Chronic Inflammatory Diseases. Front Physiol. 2021;12:730829.

8. Koivuniemi A. The biophysical properties of plasmalogens originating from their unique molecular architecture. FEBS Lett. 2017;591(18):2700–2713.

9. Jain IH, Calvo SE, Markhard AL, et al. Genetic Screen for Cell Fitness in High or Low Oxygen Highlights Mitochondrial and Lipid Metabolism. Cell. 2020;181(3):716–727 e711.

10. Yu J, Qu L, Xia Y, et al. TMEM189 negatively regulates the stability of ULK1 protein and cell autophagy. Cell Death Dis. 2022;13(4):316.

11. Perez MA, Clostio AJ, Houston IR, et al. Ether lipid deficiency disrupts lipid homeostasis leading to ferroptosis sensitivity. PLoS Genet. 2022;18(9):e1010436.

12. Zou Y, Henry WS, Ricq EL, et al. Plasticity of ether lipids promotes ferroptosis susceptibility and evasion. Nature. 2020;585(7826):603-608.

13. Cui W, Liu D, Gu W, Chu B. Peroxisome-driven ether-linked phospholipids biosynthesis is essential for ferroptosis. Cell Death Differ. 2021;28(8):2536–2551.

14. Liu J, Sun M, Sun Y, Li H. TMEM189 promotes breast cancer through inhibition of autophagy-regulated ferroptosis. Biochem Biophys Res Commun. 2022;622:37–44.

15. Lackner K, Sailer S, van Klinken JB, et al. Alterations in ether lipid metabolism and the consequences for the mouse lipidome. Biochim Biophys Acta Mol Cell Biol Lipids. 2023;1868(4):159285.

16. White JK, Gerdin AK, Karp NA, et al. Genome-wide generation and systematic phenotyping of knockout mice reveals new roles for many genes. Cell. 2013;154(2):452–464.

17. Rangholia N, Leisner TM, Holly SP. Bioactive Ether Lipids: Primordial Modulators of Cellular Signaling. Metabolites. 2021;11(1).

18. Lebrero P, Astudillo AM, Rubio JM, et al. Cellular Plasmalogen Content Does Not Influence Arachidonic Acid Levels or Distribution in Macrophages: A Role for Cytosolic Phospholipase A(2)gamma in Phospholipid Remodeling. Cells. 2019;8(8).

19. Dorninger F, Forss-Petter S, Wimmer I, Berger J. Plasmalogens, platelet-activating factor and beyond - Ether lipids in signaling and neurodegeneration. Neurobiol Dis. 2020;145:105061.

20. Westerfield M. The Zebrafish Book. A Guide for the Laboratory Use of Zebrafish Danio* (Brachydanio) rerio. Eugene, OR.: University of Oregon Press.; 2000.

21. Renshaw SA, Loynes CA, Trushell DM, Elworthy S, Ingham PW, Whyte MK. A transgenic zebrafish model of neutrophilic inflammation. Blood. 2006;108(13):3976–3978.

22. Davison JM, Akitake CM, Goll MG, et al. Transactivation from Gal4-VP16 transgenic insertions for tissue-specific cell labeling and ablation in zebrafish. Dev Biol. 2007;304(2):811–824.

23. Hall C, Flores MV, Storm T, Crosier K, Crosier P. The zebrafish lysozyme C promoter drives myeloid-specific expression in transgenic fish. BMC Dev Biol. 2007;7:42.

24. Walton EM, Cronan MR, Beerman RW, Tobin DM. The Macrophage-Specific Promoter mfap4 Allows Live, Long-Term Analysis of Macrophage Behavior during Mycobacterial Infection in Zebrafish. PLoS One. 2015;10(10):e0138949.

25. Traver D, Paw BH, Poss KD, Penberthy WT, Lin S, Zon LI. Transplantation and in vivo imaging of multilineage engraftment in zebrafish bloodless mutants. Nat Immunol. 2003;4(12):1238–1246.

26. Kanther M, Sun X, Mühlbauer M, et al. Microbial colonization induces dynamic temporal and spatial patterns of NF-κB activation in the zebrafish digestive tract. Gastroenterology. 2011;141(1):197–207.

27. Nguyen-Chi M, Phan QT, Gonzalez C, Dubremetz JF, Levraud JP, Lutfalla G. Transient infection of the zebrafish notochord with E. coli induces chronic inflammation. Dis Model Mech. 2014;7(7):871–882.

28. Nguyen-Chi M, Laplace-Builhe B, Travnickova J, et al. Identification of polarized macrophage subsets in zebrafish. Elife. 2015;4:e07288.

29. White RM, Sessa A, Burke C, et al. Transparent adult zebrafish as a tool for in vivo transplantation analysis. Cell Stem Cell. 2008;2(2):183–189.

30. Amsterdam A, Burgess S, Golling G, et al. A large-scale insertional mutagenesis screen in zebrafish. Genes Dev. 1999;13(20):2713–2724.

31. Meeker ND, Hutchinson SA, Ho L, Trede NS. Method for isolation of PCR-ready genomic DNA from zebrafish tissues. Biotechniques. 2007;43(5):610, 612, 614.

32. Liongue C, Hall CJ, O’Connell BA, Crosier P, Ward AC. Zebrafish granulocyte colony-stimulating factor receptor signaling promotes myelopoiesis and myeloid cell migration. Blood. 2009;113(11):2535–2546.

33. Kwan KM, Fujimoto E, Grabher C, et al. The Tol2kit: a multisite gateway-based construction kit for Tol2 transposon transgenesis constructs. Dev Dyn. 2007;236(11):3088–3099.

34. Pfaffl MW. A new mathematical model for relative quantification in real-time RT-PCR. Nucleic Acids Res. 2001;29(9):e45.

35. Schindelin J, Arganda-Carreras I, Frise E, et al. Fiji: an open-source platform for biological-image analysis. Nat Methods. 2012;9(7):676-682.

36. de Oliveira S, López-Muñoz A, Candel S, Pelegrín P, Calado A, Mulero V. ATP modulates acute inflammation in vivo through Duox1-derived H2O2 production and NF-kB activation. J Immunol. 2014;192:5710–5719.

37. Candel S, de Oliveira S, López-Muñoz A, et al. Tnfa signaling through Tnfr2 protects skin against oxidative stress–induced inflammation. PLoS Biol. 2014;12(5):e1001855.

38. http://ec.europa.eu/eurostat/statistics-explained/index.php/Causes_of_death_statistics.

39. Martinez-Morcillo FJ, Canton-Sandoval J, Martinez-Navarro FJ, et al. NAMPT-derived NAD+ fuels PARP1 to promote skin inflammation through parthanatos cell death. PLoS Biol. 2021;19(11):e3001455.

40. Tyrkalska SD, Candel S, Angosto D, et al. Neutrophils mediate Salmonella Typhimurium clearance through the GBP4 inflammasome-dependent production of prostaglandins. Nat Commun. 2016;7:12077.

41. Lodhi IJ, Wei X, Yin L, et al. Peroxisomal lipid synthesis regulates inflammation by sustaining neutrophil membrane phospholipid composition and viability. Cell Metab. 2015;21(1):51–64.

42. Dorninger F, Wiesinger C, Braverman NE, Forss-Petter S, Berger J. Ether lipid deficiency does not cause neutropenia or leukopenia in mice and men. Cell Metab. 2015;21(5):650–651.

43. Hasegawa T, Hall CJ, Crosier PS, et al. Transient inflammatory response mediated by interleukin-1beta is required for proper regeneration in zebrafish fin fold. Elife. 2017;6.

44. Morales RA, Allende ML. Peripheral Macrophages Promote Tissue Regeneration in Zebrafish by Fine-Tuning the Inflammatory Response. Front Immunol. 2019;10:253.

45. Nguyen-Chi M, Laplace-Builhe B, Travnickova J, et al. TNF signaling and macrophages govern fin regeneration in zebrafish larvae. Cell Death Dis. 2017;8(8):e2979.

46. Wright LC, Nouri-Sorkhabi MH, May GL, Danckwerts LS, Kuchel PW, Sorrell TC. Changes in cellular and plasma membrane phospholipid composition after lipopolysaccharide stimulation of human neutrophils, studied by 31P NMR. Eur J Biochem. 1997;243(1-2):328–335.

47. Wallner S, Grandl M, Konovalova T, et al. Monocyte to macrophage differentiation goes along with modulation of the plasmalogen pattern through transcriptional regulation. PLoS One. 2014;9(4):e94102.

48. Amunugama K, Jellinek MJ, Kilroy MP, et al. Identification of novel neutrophil very long chain plasmalogen molecular species and their myeloperoxidase mediated oxidation products in human sepsis. Redox Biol. 2021;48:102208.

