## Supplementary material for "Peds1 deficiency in zebrafish results in myeloid cell apoptosis and exacerbated inflammation": Figures S1-S3

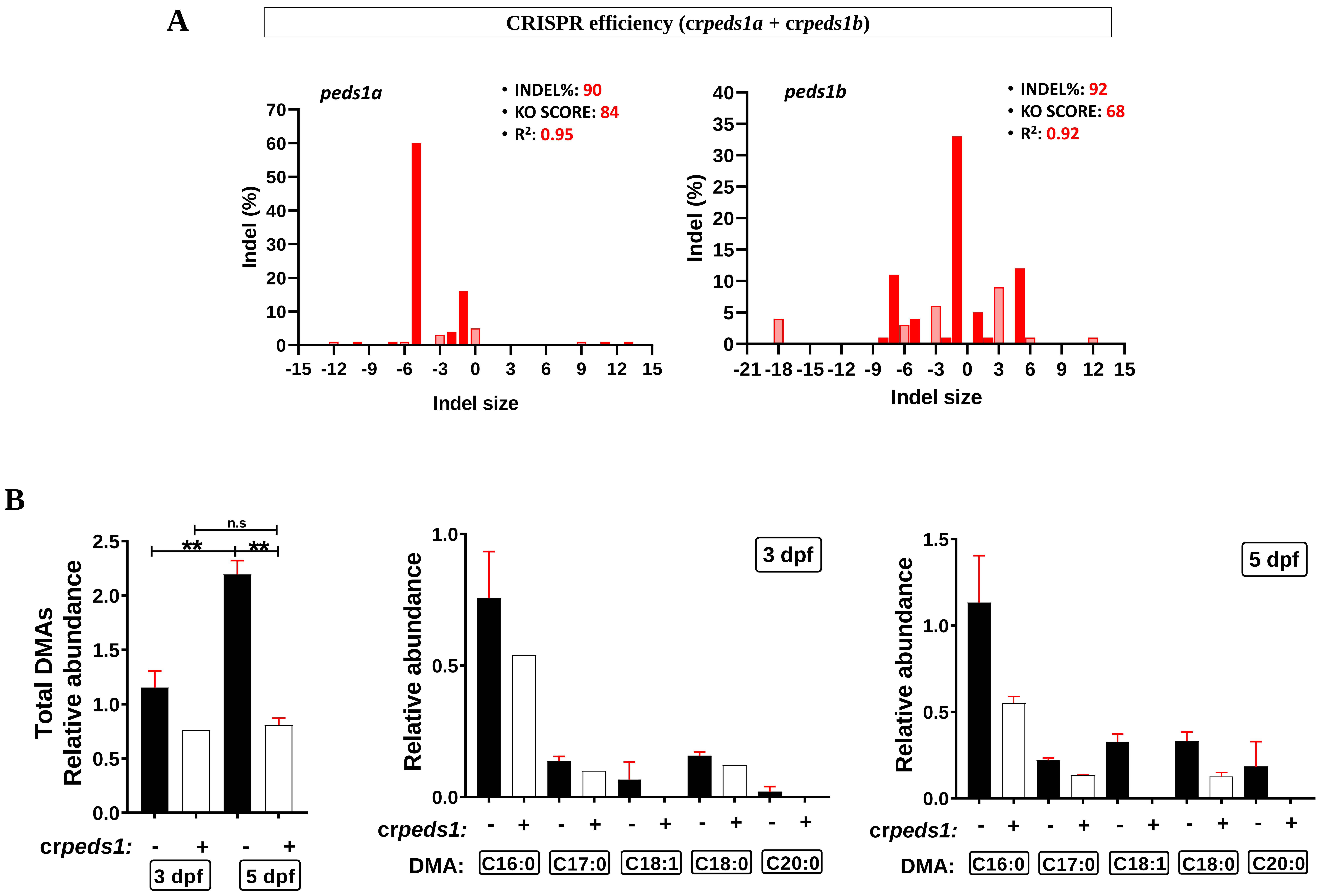


**Supplemental Figure 1 (related to Figure 1). Efficiency of *peds1* crRNA and levels of plasmalogen.** (A) Analysis of genome editing efficiency in larvae injected with *peds1a* and *peds1b* crRNA/Cas9 complexes (knock-out score 84 and 68 %, respectively) and quantification rate of non-homologous end joining-mediated repair showing all insertions and deletions (INDELS) at the target site. (B) Total and individual plasmalogen quantification at 3 and 5 dpf in cr*peds1*-injected and control larvae. The mean ± SEM of each group is shown. n.s, not significant, $**p \leq 0.01$.

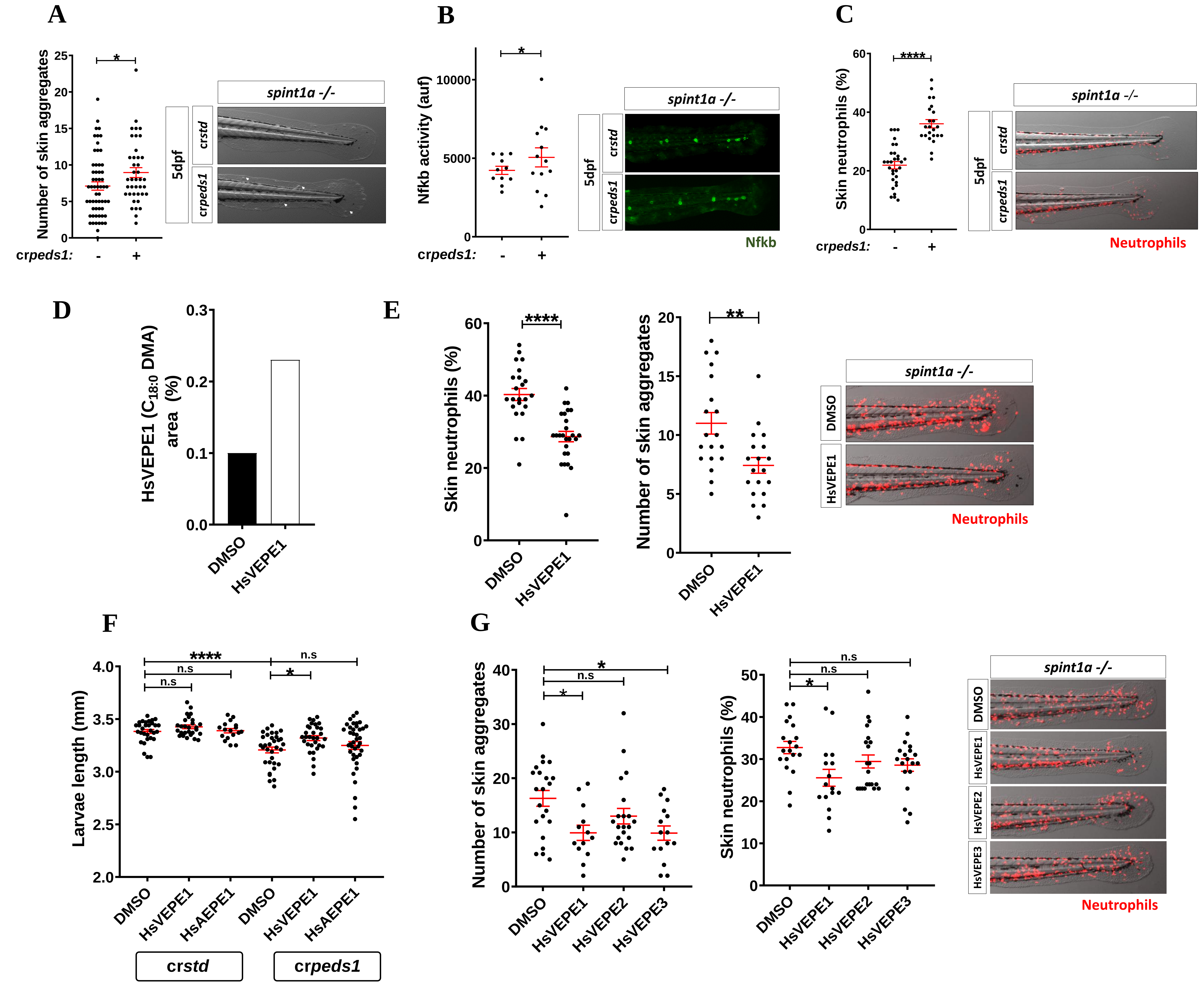


**Supplemental Figure 2 (related to Figure 2). Peds1 deficiency exacerbates skin chronic inflammation.** The same procedure represented in Figure 2A schematic was used. (A-C) Analysis and representative images of skin aggregates (white arrows) (brightfield channel), percentage of skin neutrophil (red channel; *Tg(lyz:DsRED2)*$^{nz50}$) and Nfkb activity (green channel; *Tg(NFkB-RE:eGFP)*$^{sh235}$) in cr*std* and cr*peds1* microinjected Spint1a-deficient larvae at 5 dpf. (D) Analysis of HsVEPE1 levels in Spint1a-deficient larvae treated with 20 µM of HsVEPE1 or vehicle (DMSO) for 48 hours. (E) Percentage of skin neutrophil, number of skin aggregates and representative images of Spint1a-deficient larvae treated with 20 µM of HsVEPE1 or vehicle (DMSO) for 48 hours. (F) Analysis of larval length of Spint1a-deficient larvae microinjected with cr*peds1* or cr*std* and treated with 20 µM HsVEPE1, 20 µM HsAEPE1 or vehicle (DMSO) for 48 hours. (G) Number of skin aggregates, percentage of skin neutrophil and representative merge images (brightfield and red channel; *Tg(lyz:DsRED2)*$^{nz50}$) of Spint1a-deficient larvae treated with 20 µM HsVEPE1, 20 µM HsVEPE2, 20 µM HsVEPE3 or vehicle (DMSO) for 48 hours. Each point represents one larva and the mean ± SEM of each group is also shown. *P* values were calculated using one-way ANOVA and Tukey's multiple range or unpaired Student's *t*-test. n.s, not significant, * $p \leq 0.05$, **$p \leq 0.01$, ***$p \leq 0.001$, ****$p \leq 0.001$. auf, arbitrary units of fluorescence.

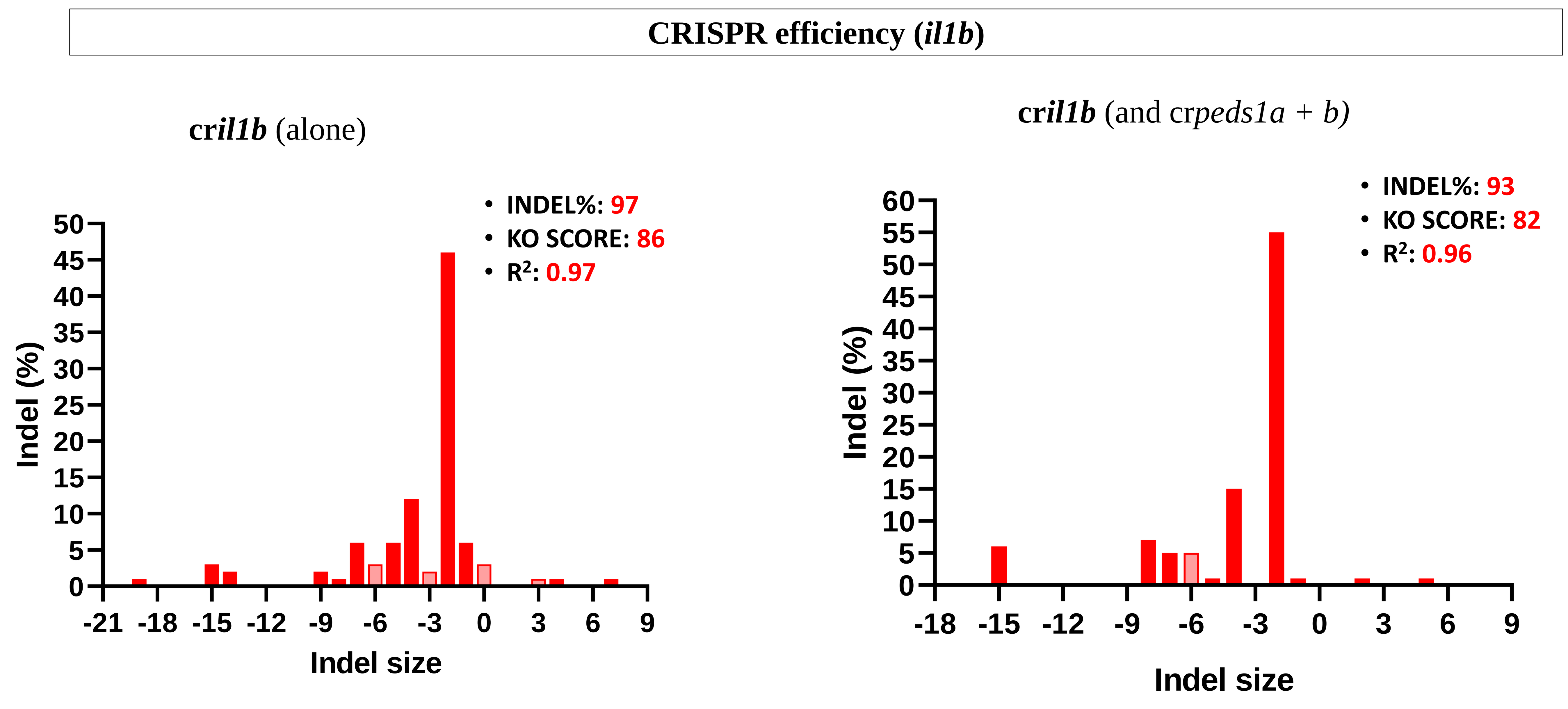


**Supplemental Figure 3 (related to Figure 4). Efficiency of the crRNA of *il1b*.** Analysis of genome editing efficiency in larvae injected with *il1b* crRNA/Cas9 complexes alone or in combination with *peds1* crRNA and quantification rate of non-homologous end joining-mediated repair showing all insertions and deletions (INDELS) at the target site.
