## Supplementary material for "Peds1 deficiency in zebrafish results in myeloid cell apoptosis and exacerbated inflammation": Table S1

**Table S1.** crRNAs and primers used in this study. The gene symbols followed the Zebrafish Nomenclature Guidelines (http://zfin.org/zf_info/nomen.html).

| **Gene** | **ENA or Ensembl accesion number** | **Name** | **Sequence (5'→3')** | **Use** |
|---|---|---|---|---|
| *peds1a* | ENSDARG00000011498 | Dr.Cas9.SI:CH211-212O1.2.1.AQ | TCCCCAATGGACCATCCCAG | CRISPR-Cas9 |
| *peds1b* | ENSDARG00000042732 | Dr.Cas9.TMEM189.1.AC | CCAGGTGGAAGTATGTTACC | |
| *il1b* | ENSDARG00000098700 | Dr.Cas9.IL1B.1.AA | CAGGCCGTCACACTGAGAGC | |
| *peds1a* | ENSDARG00000011498 | F | TGTGCCTCTCACTCTTCATTGT | Genetic edition efficiency |
| | | R | ACTGTCGACTAAGCAAAATCTCC | |
| *peds1b* | ENSDARG00000042732 | F | CACACAGTGAGACCAGCTAATCTTT | |
| | | R | CACCATACCGCCCAGGTCTA | |
| *il1b* | ENSDARG00000098700 | F | CATGATGACTTTTGTGGAGAGAAAA | |
| | | R | GTAACCTGTACCTGGCCTGC | |
| *rps11* | NM_213377.1 | F | ACAGAAATGCCCCTTCACTG | RT-qPCR |
| | | R | GCCTCTTCTCAAAACGGTTG | |
| *il1b* | NM_212844.2 | F | GCCTGTGTGTTTGGGAATCT | |
| | | R | TGATAAACCAACCGGGACA | |
| *nfkb1* | ENSDARG00000105261.2 | F | TTCTTCTTGGTCACGTGCAG | |
| | | R | ACTCTCAGCATCCGCATCTT | |
| *tnfa* | NM_212859.2 | F | GCGCTTTTCTGAATCCTACG | |
| | | R | TGCCCAGTCTGTCTCCTTCT | |
| *cxcl8a* | XM_001342570.7 | F | GTCGCTGCATTGAAACAGAA | |
| | | R | CTTAACCCATGGAGCAGAGG | |
